# Simulation-guided design of exotendons to reduce the energetic cost of running

**DOI:** 10.64898/2026.04.07.717115

**Authors:** Jon Stingel, Nicholas Bianco, Carmichael Ong, Steve Collins, Scott Delp, Jennifer Hicks

**Affiliations:** Department of Mechanical Engineering, Stanford University, Stanford, California, United States of America; Department of Bioengineering, Stanford University, Stanford, California, United States of America

**Author notes:** Corresponding author: (JH).

## Abstract

A passive device that attaches to the feet, called an exotendon, can reduce the energetic cost of running at moderate speeds, but its efficacy and optimal design parameters at higher speeds are unknown. Identifying optimal parameters at new speeds experimentally would require many experimental trials with different exotendon designs, which is challenging for participants at higher running speeds. We developed a muscle-driven simulation framework to predict the effect of various exotendon designs on the energetic cost of running at an experimentally untested speed (4 m/s). We used these predictions to select four designs, which we evaluated experimentally as users ran at this speed. The framework correctly predicted that an exotendon that reduced energetic cost at 2.7 m/s would also reduce energetic cost at 4 m/s (10% predicted vs. 5.7% measured) and that a short, stiff exotendon and a long, compliant exotendon would not significantly reduce energetic cost. However, exotendon parameters predicted by the simulation to maximize energetic savings did not significantly reduce energetic cost when evaluated experimentally. There was variability between participants in both the magnitude of maximum energy savings and the exotendon condition associated with those savings. In a 5-km time trial performed with and without the exotendon condition that elicited the largest energy savings for each participant during the experiment, we observed a lower average heart rate (-3.9 ± 3.8 beats/min; *P*=0.03; mean ± standard deviation) and increased cadence (15.9 ± 9.6 steps/min; *P*=0.002) when participants ran with the exotendon but did not observe a statistically significant difference in finishing time (-13.5 ± 24.6 sec; *P*=0.3). These results demonstrate exotendons can reduce energetic cost across multiple running speeds and that predictive simulations provide a framework for guiding experiments to evaluate assistive device designs.

**Author summary:** Designing assistive devices that help people move more efficiently usually requires many experimental trials. These studies can be time-consuming and physically demanding, especially when testing multiple device designs. In this study, we explored whether computer simulations could help guide the design of an assistive device for running called an exotendon. The exotendon is a simple elastic band that connects the feet and can help runners use less energy. Previous experiments showed that the device reduces the energy needed to run at moderate speeds, but it was unclear whether it would also work at faster speeds or which design would lead to energetic savings. We first used simulations of human running to test many possible exotendon designs at a faster speed. These simulations allowed us to identify promising designs before conducting experiments. We then tested a small number of these designs with runners. The experiments confirmed that the exotendon can reduce the energy required to run at faster speeds, although the efficacy of different designs varied between individuals. Our results show that computer simulations can help researchers rapidly evaluate a variety of assistive device ideas and focus experimental testing on the most promising designs.

## I. Introduction

Wearable assistive devices are being developed for sports applications, where even modest improvements in energetic efficiency can translate to meaningful performance gains. In running, much of this effort has focused on footwear, where advances in shoe geometry and materials have improved running economy and speed, contributing to record-setting performances (1–9). Beyond footwear, several studies have demonstrated energetic savings for both active and passive devices during running. These studies employ different assistance strategies, including active ankle moment assistance (10), passive moments about the hip (11), and assistance via the exotendon—a device that connects a runner’s shoes via an extension spring (12).

The exotendon is a simple, passive device that has been shown to reduce the energetic cost of running at 2.7 m/s by 6-8% (12,13). It functions as an extension spring connecting a runner’s shoes and has two design parameters: the stiffness and slack length of the spring. Runners wearing the exotendon tend to increase their cadence and reduce their hip and knee moments (12). These biomechanical changes decrease demand on the quadriceps and hip flexors, leading to lower energetic cost (13). Such savings could allow runners to increase their speed without increasing energetic cost, but it remains unknown whether the benefits of the exotendon are present at faster running speeds. Moreover, exotendon design parameters have not been systematically evaluated across speeds and may require re-tuning to achieve benefits during fast running.

Designing and evaluating assistive devices is challenging because performance benefits depend on each design parameter (1,2,14). Further, even seemingly simple interventions, such as footwear, can elicit complex and unexpected biomechanical responses. As the number of adjustable design parameters increases, experimental evaluation becomes impractical due to the time and physical demands placed on participants. These challenges are amplified in high-performance running, where measuring energetic cost requires several minutes of steady-state locomotion below the participant’s aerobic threshold (15). Musculoskeletal simulations could streamline the evaluation of assistive technologies by enabling rapid, systematic screening of candidate designs, thus identifying a narrower parameter space where runners are likely to benefit without the need for exhaustive physical testing (16–19). Making accurate predictions in simulation remains a challenging task, however, particularly in the presence of assistive devices (20,21).

The goal of this work was to design and evaluate the effectiveness of exotendons while running at 4 m/s via predictive simulations coupled with targeted experimental testing. This speed was chosen to be markedly faster than the previously tested 2.7 m/s, while still falling in a range that can be maintained by semi-elite runners for the duration of a marathon (22). We first developed musculoskeletal simulations to predict the energetic effect of exotendons across a range of stiffness and slack-length combinations while running at 4 m/s. We used these predictions to select four configurations for experimental testing and preregistered our results (23). We then measured runners’ energetic cost while using each design and evaluated their performance with their individually best-performing design during a 5-km time trial. This paired simulation-experimental framework allowed us to streamline device testing and assess the simulation’s accuracy in selecting device parameters. Based on our simulations, we expected the exotendon to provide energetic benefits at the faster running speed.

## II. Results

### ***A.*** Simulation results

We developed a muscle-driven simulation framework to estimate the energetic cost of a running gait cycle with an exotendon. The framework was implemented with OpenSim Moco (24) and takes as input a generic musculoskeletal model, the desired running speed, a set of experimentally measured, unassisted (referred to as natural) joint coordinates and ground reaction forces, and slack length and stiffness values for the exotendon. The framework finds an optimized set of joint coordinates and ground reaction forces, along with the muscle activations, that minimize energetic cost given the provided exotendon parameters (see Methods, Section IV.B). The simulation framework reproduced changes in experimental energetic cost (mean error: 3.0%; 95% CI: -0.3 to 6.2%) (Fig S1), joint kinematics (root-mean-squared-error (RMSE): 4.8 ± 2.2°; Pearson *r* = 0.97 ± 0.01; mean ± standard deviation), and joint moments (RMSE: 0.3 ± 0.1 Nm/kg; *r* = 0.94 ± 0.02) at the hip, knee, and ankle during running at 2.7 m/s with the previously published exotendon design (Fig S2) (12,13). We verified that the simulation framework reproduced experimental joint kinematics (RMSE: 8.4 ± 4.2 degrees; *r* = 0.94 ± 0.03) and moments (RMSE: 0.5 ± 0.1 Nm/kg; *r* = 0.91 ± 0.04) at the hip, knee, and ankle in natural running at 4 m/s (25) (Fig S4).

We used this predictive simulation framework to estimate the effects of 25 combinations of exotendon slack length (*l*) and stiffness (*k*) on the energetic cost of running at 4 m/s for a generic musculoskeletal model (Table 1). The framework predicted reductions in energetic cost compared to simulated natural running across most designs, with relatively small differences between many of the top-performing designs. The previously published exotendon design (“medium-original” in Table 1), used a medium length (*l*=0.29) and medium stiffness (*k*=120). For this design, the simulation predicted a 10% reduced energetic cost. The largest predicted reduction (12%) occurred with a long-stiff exotendon (*l*=0.43, *k*=240).

**Table 1.**
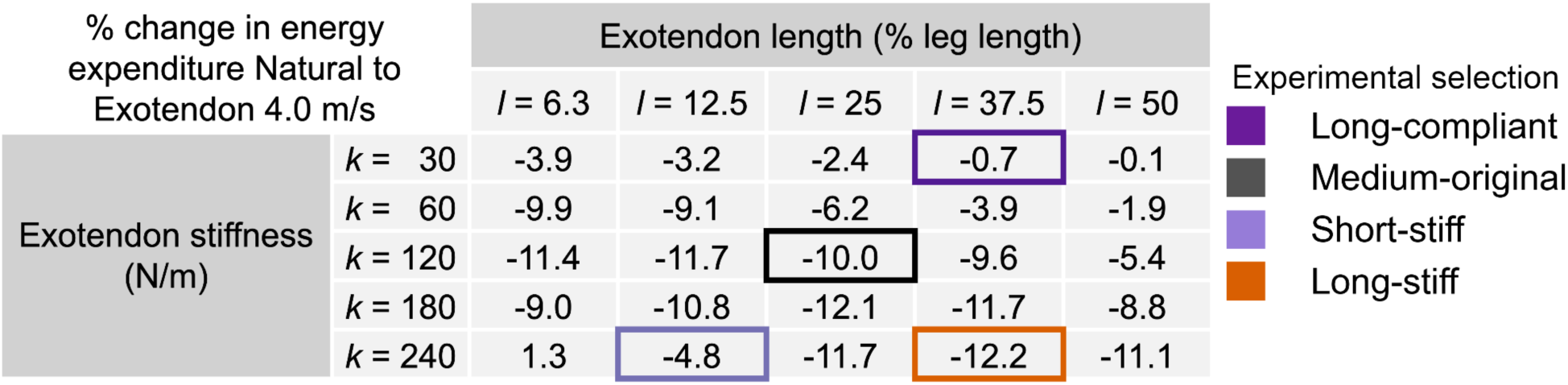
Predicted effect of simulated exotendon designs on the energetic cost of running at 4.0 m/s. Values are the percent change in average energetic cost (W/kg) relative to natural running for each simulated combination of exotendon stiffness (*k*) and length (*l*). Negative values indicate lower energetic cost for running with an exotendon compared to natural, while positive values indicate an increase in energetic cost with the exotendon. Stiffness values (rows) ranged from 30 to 240 N/m, and length values (columns) ranged from 6.25-50% of leg length. Leg length in the simulation was 0.92 m. Experimentally tested designs are outlined and color-coded: long-compliant (*l*=37.5, *k*=30; dark purple), medium-original (*l*=25, *k*=120; black), short-stiff (*l*=12.5, *k*=240; light purple), and long-stiff (*l*=37.5, *k*=240; orange).

Based on the *a priori* simulation predictions, we selected four exotendon designs with a range of predicted effects for experimental testing: medium-original (*l*=25, *k*=120; 10% predicted reduction), long-stiff (*l*=37.5, *k*=240; 12%), short-stiff (*l*=12.5, *k*=240; 4.8%), and long-compliant (*l*=37.5, *k*=30; 0.7%). Prior to experiments, we preregistered our simulation results on bioRxiv (23).

### ***B.*** In-lab experimental results

We recruited eleven participants to experimentally evaluate the effects of the four exotendon designs identified in simulation (see Methods, Section IV.D). Participants trained at 4 m/s on the treadmill unassisted and with each of the designs for 6 minutes. They then completed an additional 30 minutes of training with each of the four exotendons during at-home runs. Following at-home training, participants ran on an instrumented treadmill at 4 m/s for 6 minutes using each of the exotendons and in a natural condition, with a randomized order of conditions. We used indirect calorimetry to compute participants’ energetic cost, averaged over the final minute of each trial (Fig 1) (26). Three trials across all subjects were discarded due to runners’ respiratory exchange ratio exceeding 1.0, indicating they may have exceeded their anaerobic threshold.

**Fig 1.**
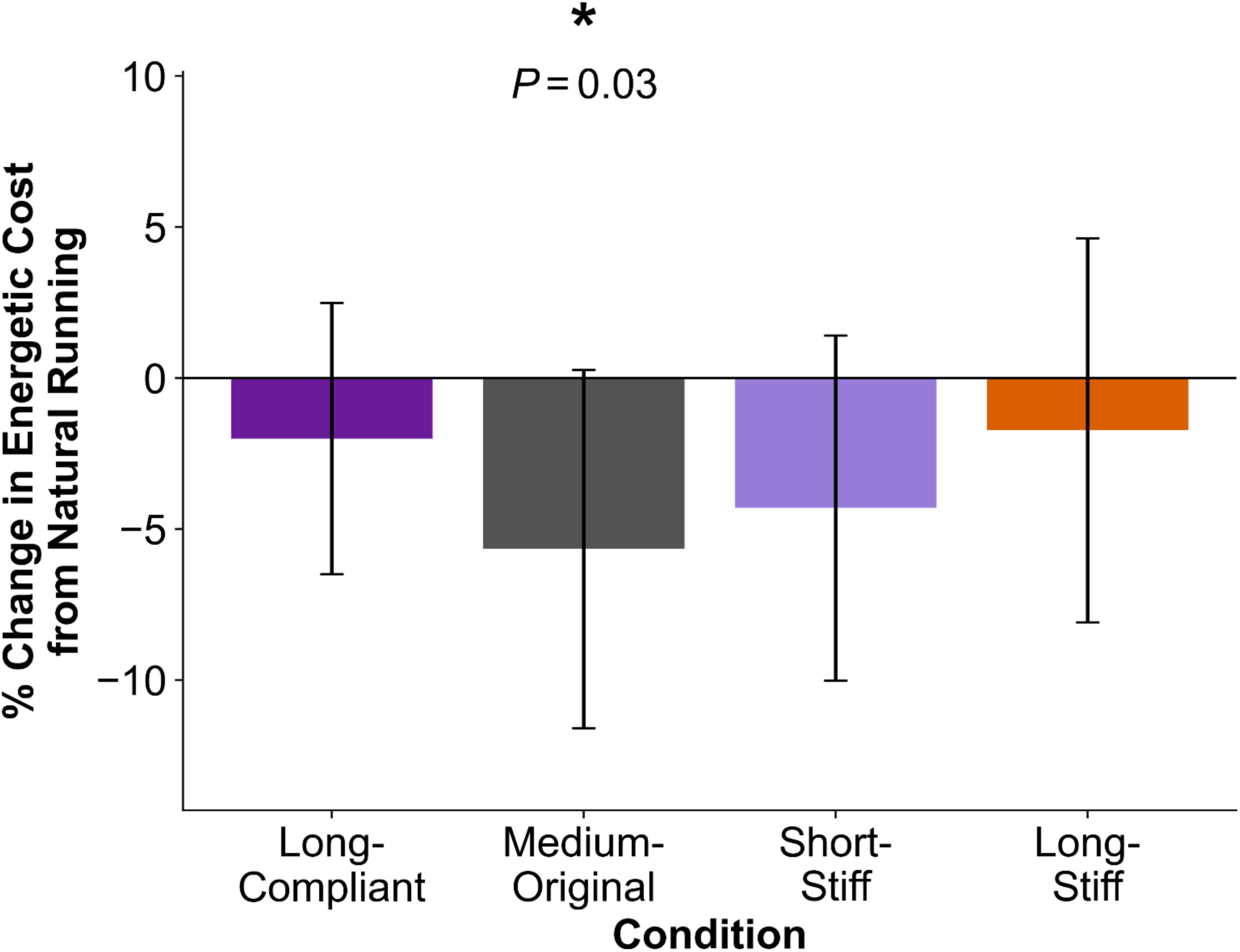
Percent change (mean ± SD) in energetic cost (net metabolic power in W/kg) relative to natural running for each exotendon condition from *N*=11 participants running at 4 m/s. Negative values indicate that the energetic cost of running was lower with the exotendon. * indicates a significant difference from natural running (4 comparison Bonferroni-corrected, α=0.05).

When running with the medium-original exotendon design, participants decreased their energetic cost by 5.7% (-0.9 ± 0.9 W/kg; *P*=0.03). There were no statistically significant changes in energetic cost for the long-compliant (-0.3 ± 0.7 W/kg; *P*=0.6), short-stiff (-0.6 ± 0.8 W/kg; *P*=0.2), or long-stiff (-0.3 ± 0.9 W/kg; *P*=1.0) designs. Participants exhibited their largest energetic cost reduction during the experiment with different designs: two runners experienced their largest reduction with the long-compliant design, six with the medium-original design, two with the short-stiff design, and one with the long-stiff design (Fig S8).

To investigate the difference between predicted and observed energetic cost changes across exotendon designs, we computed the tension forces applied by the exotendons using measured slack length, material stiffness, and exotendon stretch estimated from motion-capture markers during running. We compared experimental tension forces with those predicted by the simulation for each exotendon design (Fig 2). In both the experiment and the simulation, the short-stiff design produced the largest peak tension (179 N experimentally vs. 160 N in simulation) and the long-compliant design produced the smallest peak tension (12 N vs. 14 N). While the simulations predicted higher peak tensions for the long-stiff design (105 N) compared to the medium-original design (73 N), the two designs produced similar, intermediate peak tensions experimentally (94 N and 98 N, respectively).

**Fig 2:**
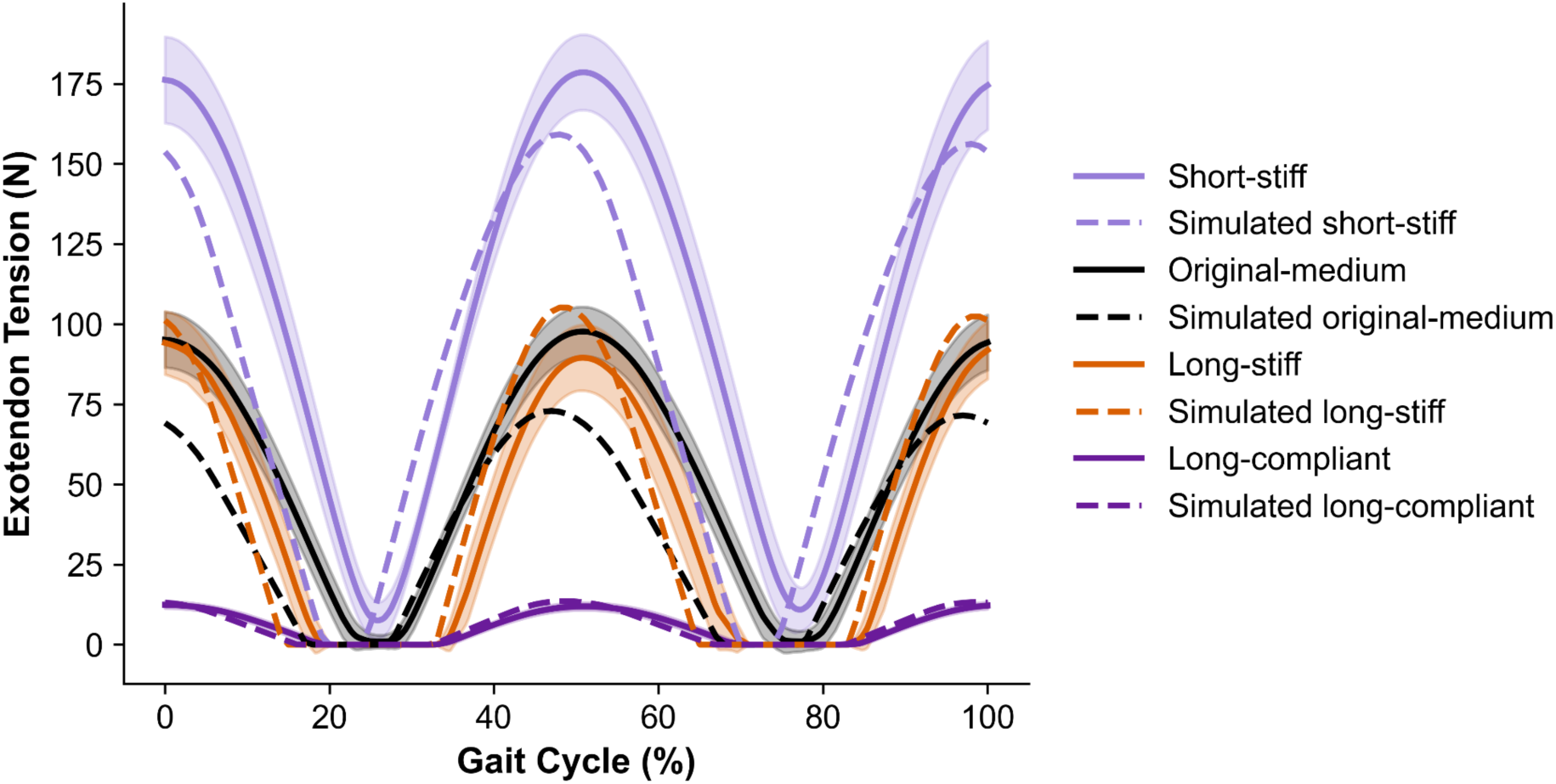
Experimental and simulated exotendon tension across the gait cycle. For the experiments, we computed the mean tension across participants (solid line) and standard deviation (shaded region) from measured material stiffness, customized slack length, and the distance between foot attachment points, estimated from motion capture markers. Corresponding simulated tension profiles are shown for comparison (dotted lines). Values are plotted over a full gait cycle, beginning at heel strike.

### ***C.*** Out-of-lab experimental results

To determine if the energetic savings from using the exotendon translated into increased running speed, ten runners completed two 5-km time trials (see Methods, Section IV.D). The first run occurred on the first day of the experimental protocol and was unassisted. In the second 5-km trial, which occurred on the last day of the protocol, participants ran with the exotendon that elicited the largest reduction in energetic cost during the treadmill testing sessions. One runner was unable to complete their final 5-km trial, leaving *N*=10. Of the 10 remaining runners, 8 reduced their 5-km run time. All runners who used the medium-original and long-compliant exotendons improved their time, whereas those who used the long-stiff and short-stiff exotendon increased their run time. On average, however, we did not detect a difference in 5-km time between conditions (-13.5 ± 24.6 seconds; *P*=0.3), due to the large variability between participants; one runner increased their time by 31 seconds, while another decreased time by 67 seconds. Average heart rate decreased when running with the exotendon (-3.9 ± 3.8 bpm; *P*=0.03) and cadence increased (15.9 ± 9.6 spm; *P*=0.002) (Fig 3). Average run speed during the natural and exotendon 5-km runs was 4.5 ± 0.2 m/s, which was faster than during treadmill testing.

**Fig 3.**
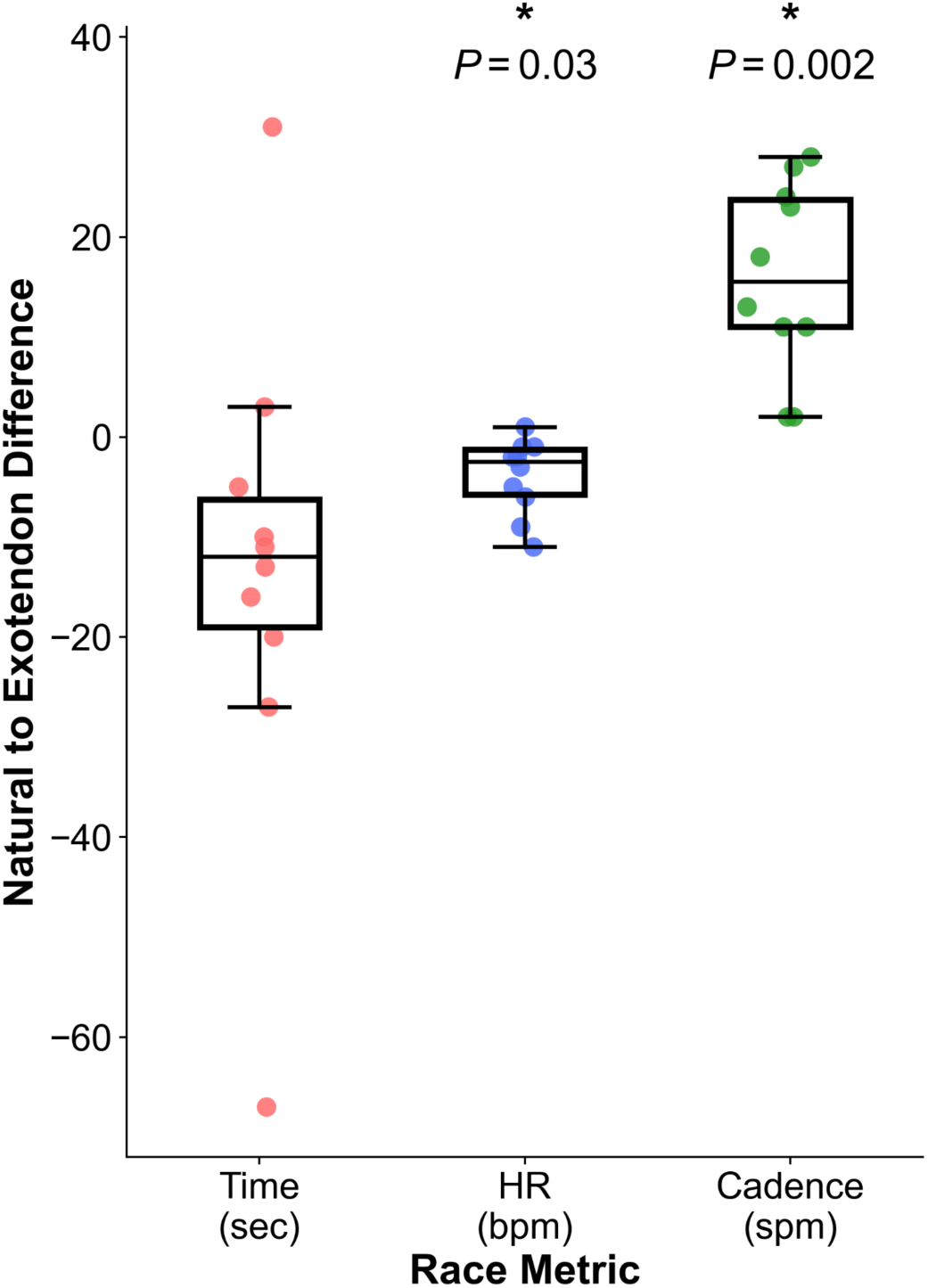
Changes in 5-km performance measures with the best-performing exotendon for each runner. Differences between exotendon and natural conditions are shown for 5-km time (red), average heart rate (blue), and step cadence (green). * indicates a significant difference between conditions (3 comparison Bonferroni-corrected, α=0.05).

## III. Discussion

The goal of this study was to evaluate the use of musculoskeletal simulations to streamline the design and experimental evaluation of exotendons for running. Simulations indicated that exotendons would reduce energetic cost when running at 4 m/s across a range of stiffness–length combinations, with both the medium-original exotendon design and a longer, stiffer configuration predicted to provide substantial benefit. Experimental testing confirmed that exotendons reduced energetic cost at 4 m/s, although the energetically optimal design varied between participants. These results demonstrate that a simple, passive assistive device can reduce energetic cost across multiple running speeds. The original exotendon design reduced energetic cost by approximately 6-8% at 2.7 m/s (12,13) and by 5.7% at 4 m/s in the present study.

Together, these findings suggest that the exotendon’s ability to reduce energetic cost may generalize beyond a single speed, expanding the range of runners and contexts in which it may provide energetic benefit.

This work demonstrates the utility of predictive simulations as a tool for assistive device design. The framework enabled evaluation of 25 exotendon configurations without exhaustive experimental testing, reducing participant burden by more than 14 hours of running time per individual. The simulated exotendons that elicited the largest reductions in energetic cost had an average peak tension of ∼100 N, consistent with the medium-original exotendon (98 N) that produced significant cost reductions experimentally. The long-stiff exotendon was predicted to generate 105 N of tension and produced 94 N on average experimentally; however, this design did not yield a statistically significant reduction in energetic cost, contrary to simulation predictions. Simulations predicted that many exotendon designs would provide substantial energetic cost savings, with nine configurations reducing cost by 10-12%. Given that typical measurement variability in indirect calorimetry is on the order of 2-3% (26), our ability to experimentally resolve differences between similarly performing designs is limited. Importantly, simulations correctly predicted that exotendons would reduce energetic cost at faster running speeds and were used to narrow the design space in advance of experimental testing, enabling a feasible and targeted protocol. This approach could provide a framework for integrating predictive simulation with experimental validation of wearable assistive devices.

While the simulations showed utility in guiding the subsequent experiments, the experimental testing of our simulation-driven hypotheses also uncovered limitations of the simulation framework that could be improved in future work. Our simulation framework relied on several modeling assumptions that may have influenced how accurately predicted exotendon responses reflected human coordination strategies. We designed the cost function to minimize muscle effort, a common approach that accurately recreates gait dynamics in simulation (27).

When individuals are running and utilizing an assistive device, other factors (e.g. stability or comfort) may influence their gait coordination strategy. These factors were not captured in the current optimization. Furthermore, our framework included several simplifications: the musculature in the upper limbs and torso of the model were not included, and we included only sagittal plane degrees of freedom. Prior studies, however, have shown that the majority of kinetic and energetic changes with the exotendon occur in the lower extremities for muscle groups that actuate sagittal plane degrees of freedom (12,13).

On average, the medium-original design led to the largest energetic cost savings experimentally; however, not all runners achieved their minimal energetic cost with this design, and at least one runner exhibited their minimum with each of the four tested exotendons. We cannot confirm that individuals consistently benefit from a personalized design, as we did not perform an independent validation trial using each runner’s observed best-performing exotendon. Furthermore, given the limited sample size and typical variability in indirect calorimetry measurements, differences between designs may fall within measurement error, limiting our ability to distinguish between them. This inter-individual variability nonetheless underscores the importance of exploring personalization in assistive device design and suggests that subject-specific factors—such as anthropometrics, natural running kinematics (e.g., stride length), training history, adaptation rate, or preference—may influence how effectively assistance is utilized. Future simulation-based design frameworks could address this by simulating a personalized version of the model (rather than a generic model, as we used).

Average heart rate decreased during the time trial despite no detectable change in time performance, suggesting reduced physiological effort without a measurable improvement in speed. The absence of time improvements may reflect a mismatch between the treadmill training and testing speed (4 m/s) and the self-selected race pace during the time trial (4.5 ± 0.2 m/s).

Since the runners did not train at their 5-km pace, they may not have been able to utilize the exotendon effectively at that speed. Performance changes during the 5-km run were variable, with one runner increasing their time by 31 seconds and another decreasing by 67 seconds.

Notably, all runners who completed the time trial with the medium-original and long-compliant exotendons reduced their run time, whereas those that used the short-stiff and long-stiff exotendons did not. Given that the medium-original exotendon also produced significant energetic savings, these observations suggest that focusing on this design may have yielded more consistent performance benefits. However, given the small sample size of the current study, these findings should be interpreted cautiously. Future studies should evaluate exotendon effects across a wider range of running speeds and incorporate structured familiarization at race-relevant paces to determine whether energetic savings can translate to improved time.

In our experiment, we attempted to provide runners sufficient training time to adapt to each exotendon. While we saw energetic reductions, it is possible that the runners were still adapting. Studies have shown that continued task training leads to even greater savings (28), and our runners may have benefited from longer training. A further limitation of our experiment was our small and majority-male cohort.

The exotendon is a simple passive device capable of reducing energetic cost across multiple running speeds. This study demonstrates that its benefits can extend to faster running, while highlighting the role of predictive musculoskeletal simulation in guiding device design. More broadly, this integrated framework of simulation-based design and experimental validation provides a valuable approach for accelerating the development and refinement of wearable assistive technologies.

## IV. Materials and Methods

### ***A.*** Musculoskeletal model

We used a sagittal-plane model with 18 muscle actuators to represent the major muscle groups that drive sagittal-plane motion in order to prioritize computational efficiency and enable rapid evaluation of exotendon parameter variations while maintaining biofidelity (Fig 4). The skeleton, which was based on models from prior simulation studies (18, 29–31), had 20 degrees-of-freedom (DOFs) and was implemented in OpenSim 4.5. The model included a pelvis-ground joint with three rotational and two translational DOFs; sagittal-plane joints with one rotational DOF at the hip, knee, ankle, metatarsophalangeal joint, shoulder, and elbow; and a lumbar joint with three rotational DOFs. We scaled the generic model to match the anthropometric data of a representative runner (male, age: 27, height: 178 cm, mass: 73 kg) from a previously collected dataset (13).

**Fig 4.**
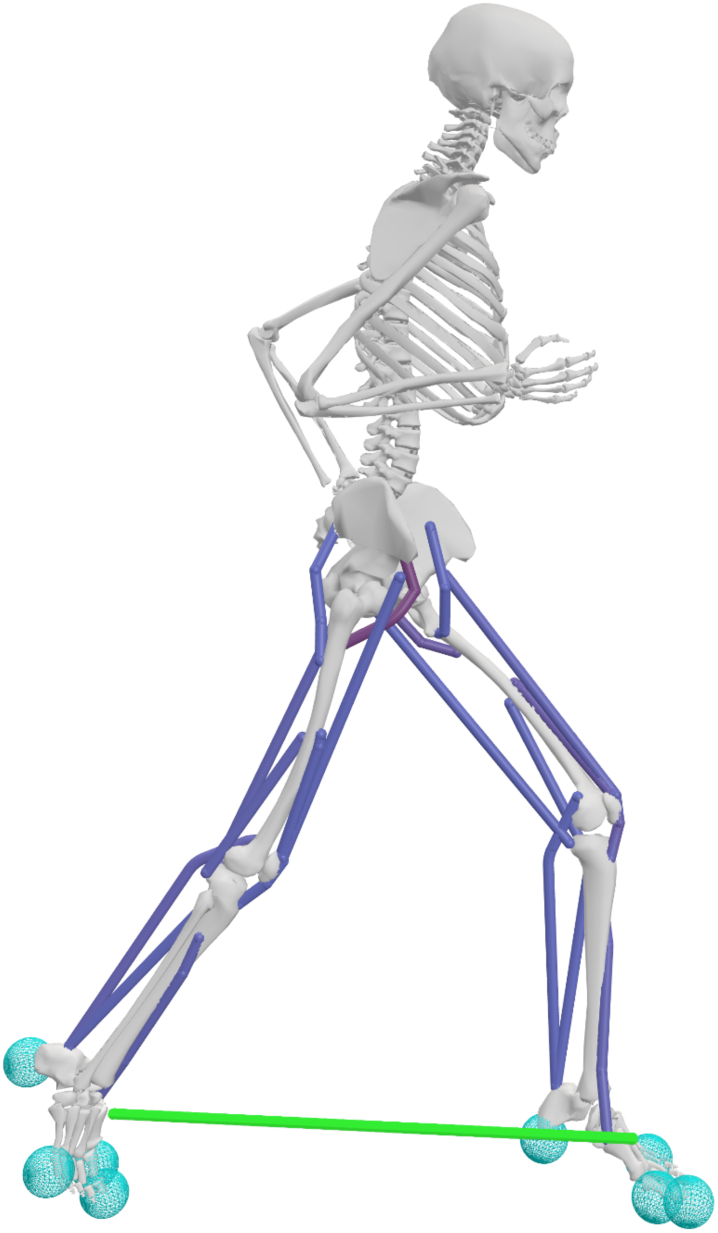
OpenSim musculoskeletal model used for predictive simulations. The model utilizes 18 Hill-type muscles that actuate the lower limbs (dark blue), and the exotendon is modeled as a linear spring (green). Foot-ground contact was modeled in the feet using eight Hunt-Crossley contact spheres (light blue) (32).

Eighteen Hill-type muscle-tendon units actuated the lower extremities, and idealized torque generators actuated the upper limb and lumbar joints (33,34). Nine muscle groups were modeled per leg: iliopsoas, gluteus maximus, rectus femoris, biarticular hamstrings, vasti, biceps femoris short head, gastrocnemius, soleus, and tibialis anterior (35,36). To approximate the force-generating capacity of trained runners and ensure sufficient torque production across joint ranges of motion, we doubled maximum isometric muscle forces and increased the active force-length operating range by 50% (37–40). Passive force elements at the lumbar and metatarsophalangeal joints represented force contributions from ligaments and other soft tissues (37,41).

We modeled foot-ground contact using four smoothed Hunt–Crossley contact spheres in each foot (32). One sphere was located in the heel, one in the forefoot, and two at the metatarsophalangeal joint in the rearfoot. All contact spheres had the following parameters: stiffness of 10 MPa, dissipation coefficient of 1.0 s/m, static and dynamic friction coefficients of 0.8, viscous friction coefficient of 0.5, transition velocity of 0.2 m/s, and radius of 0.035 m.

We modeled the exotendon as a linear extension spring connecting points on the calcanei (as measured with motion capture in prior studies), parameterized by slack length and spring stiffness. To evaluate the effects of these parameters on the energetic cost of running, we tested slack lengths of 6.25, 12.5, 25, 37.5, and 50% of leg length and stiffness values of 30, 60, 120, 180, and 240 N/m. A slack length 25% of leg length and stiffness of 120 N/m corresponds to the previously published exotendon (12). We simulated each combination of stiffness and slack length, resulting in a total of 25 model variants.

### ***B.*** Simulation framework

We generated muscle-driven tracking simulations with OpenSim Moco (24) to evaluate the energetic effects of candidate exotendon designs. To generate simulations, we used previously published treadmill running datasets at 2.7 and 4 m/s on a treadmill (13,25). For each speed, we selected three-dimensional (3D) motion capture and ground reaction force data from a representative subject over a single gait cycle of natural running. We computed joint coordinate trajectories using OpenSim’s Inverse Kinematics tool (42), and used the resulting kinematics and measured ground reaction forces as reference data for simulations at the corresponding speeds.

Simulation inputs included the reference inverse kinematic joint coordinate trajectories 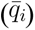, 3D ground reaction forces 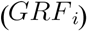, desired running speed, initial and final times (*t_i_* and *t_f_*, respectively), and the musculoskeletal model. For each exotendon simulation, we updated the corresponding spring parameters in the model.

Moco solved for muscle activations (*a_i_*), joint kinematics (*q_i_*), and ground reaction forces (*GRF_i_*) that reproduced motion consistent with the tracked reference data, and minimized the objective function (Eq. 1):

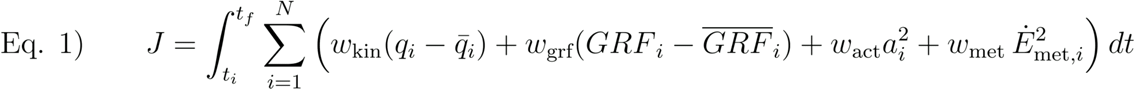

where 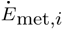 is total muscle metabolic rate. In Eq. 1, the kinematic tracking (*w*_kin_ = 1.0 × 10^−6^), contact tracking (*w_grf_* = 14), activation (*w_act_* = 8.0), and metabolics cost (*w*_met_ = 4.0 × 10^−5^) weights were selected to balance contributions across terms such that activations and metabolics constituted the majority of the objective value. This balance was selected to ensure that muscle effort terms (activation and metabolic cost) were larger.

To improve computational efficiency, we performed simulations over half a gait cycle with symmetry enforced across lower limb kinematics, muscle activations, and tendon forces.

We first simulated natural and exotendon-assisted running at 2.7 m/s to verify that the framework reproduced previously reported kinematic, kinetic, and energetic changes when running with an exotendon. We compared differences in joint kinematics and kinetics between simulated natural and exotendon running to published data using RMSE and Pearson correlation (12,13). We also simulated natural running at 4 m/s and computed differences in joint kinematics and kinetics to published results using RMSE and Pearson correlation (25).

We then simulated exotendon-assisted running at 4 m/s across all 25 combinations of slack length and stiffness. Average energetic cost was computed for each condition over the gait cycle. Because exotendons can alter the energetically optimal stride duration, we simulated four stride durations for each design and selected the duration that minimized energetic cost in each design for subsequent analyses (Fig S9). The tested stride durations (90%, 95%, 100%, and 105%) were defined relative to the reference runner’s natural stride duration from the previous experiment (13). We computed exotendon force profiles from spring stiffness and stretch during the energetically optimal stride. All code and supporting materials are available in an online repository: https://github.com/stingjp/ExotendonPredictions.git.

Based on the simulation results, we selected four exotendon configurations for experimental testing: the previously published design (medium-original), the predicted optimal design with longer slack length and greater stiffness (long-stiff), a shorter and stiffer configuration (short-stiff), and a longer and more compliant configuration (long-compliant). We preregistered these selections and the predicted energetic savings prior to the experimental evaluation (23).

### ***C.*** Participants in experiment

Eleven healthy adults (1 female, 10 males; age: 28 ± 6 years; height: 177 ± 8 cm; mass: 68 ± 7 kg; mean ± SD) participated in the study. Participants were at least 18 years old, ran a minimum of three times per week, had no running-related injuries in the prior six months, and could complete a 5-km race in under 20 minutes. The study was approved by the Stanford University Institutional Review Board, and all participants provided written informed consent prior to participation.

### ***D.*** Experimental protocol

The experimental protocol consisted of four laboratory visits per participant over approximately two weeks, with additional at-home training between the second and third visit (Fig 5). During the first visit, participants completed a maximal-effort 5-km run on the Stanford track following a self-selected warm-up. Runners were instructed to arrive rested. Run time, heart rate, and cadence were recorded using a heart rate monitor and activity watch (HRM3-SS; Forerunner 230; Garmin, Olathe, KS, USA).

**Fig 5.**
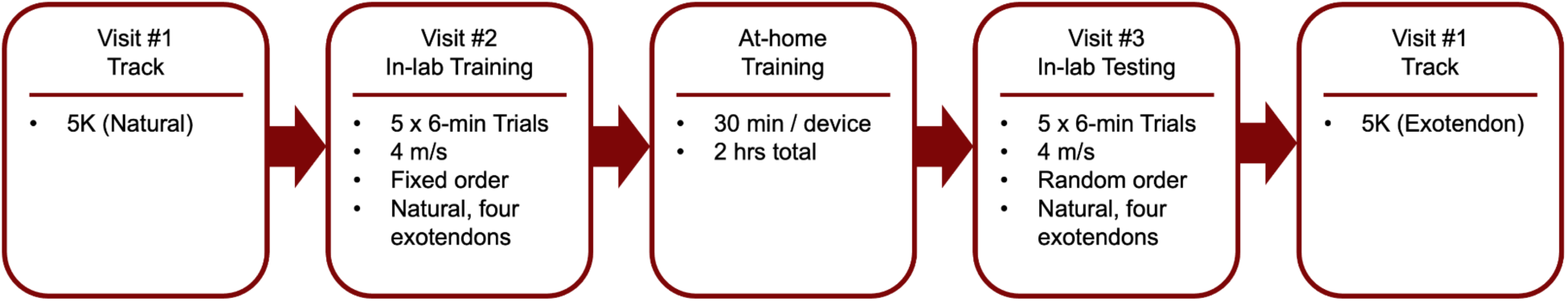
Experimental protocol covering four visits and at-home training over approximately 2 weeks. The experimental protocol outlines the order of experimental visits that runners followed, and the running tasks completed.

The second visit, conducted within 2 days of the first, introduced participants to the exotendon designs and included in-lab training runs. Participants completed five 6-minute treadmill trials (natural running and four exotendon conditions) with at least 5 minutes of rest between trials. Running speed was ramped from 3 to 4 m/s during the first minute of the trial.

Training trials were performed in a fixed order (natural, long-compliant, medium-original, short-stiff, long-stiff) beginning with the lowest peak exotendon tension.

Following this session, we provided participants all four exotendon configurations for at-home familiarization. Over approximately 1.5 weeks, they completed 30 minutes of self-selected pace running with each exotendon (2 hours total) to allow sufficient adaptation prior to primary data collection.

The third visit served as the primary in-lab data collection. Participants first completed a 5-minute standing trial to establish baseline metabolic rate using a breath-by-breath indirect calorimetry system (Quark CPET, COSMED, Rome, Italy), with energetic cost averaged over the final minute (26). We used one retroreflective marker on each shoe to measure 3D positions of the exotendon attachment points throughout the duration of each running trial. Marker positions were recorded at 200 Hz using a 10-camera optical motion capture system (Motion Analysis Corporation, Santa Rosa, CA, USA). Energetic and kinematic data was collected during five randomized 6-minute trials (natural running and four exotendon conditions) at 4 m/s, with speed ramped from 3 to 4 m/s during the first minute and 5 minutes of rest between trials.

During the final visit, participants completed a second maximal 5-km run on the Stanford track while wearing the exotendon that elicited the largest reduction in energetic cost during their third visit. This run was scheduled within 3 days of the third visit at a similar time of day as their first 5-km run. We recorded run time, heart rate, and cadence, as in the first session.

### ***E.*** Experimental data processing

We calculated experimental energetic cost from indirect calorimetry measurements by averaging the rate of energy expenditure over the final minute of each trial (26) and subtracting the static standing baseline to obtain net energetic cost. Any trials where the runner had an average respiratory exchange ratio larger than 1.0 were excluded from analysis. For each exotendon condition, percent change in energetic cost was computed relative to each runner’s natural running trial.

Exotendon tension was estimated from experimental motion capture data using the known stiffness and slack length of each design. We calculated instantaneous tension as the product of the exotendon stiffness and stretch, where stretch was determined from the distance between motion capture markers placed at the exotendon attachment points on the feet. The Euclidean distance was computed from the 3D positions of the two markers, and stretch determined by subtracting the known slack length of each exotendon. We used a minimum of five gait cycles per condition were analyzed for each runner. Tension values were first averaged across gait cycles within each runner and then across runners for each condition.

### ***F.*** Statistical analyses

Statistical analyses were performed to evaluate differences in energetic cost between exotendon conditions and natural running. We assessed normality of within-subject differences using the Shapiro-Wilk test. Then, we performed paired t-tests, comparing each of the four exotendon conditions to natural running. Bonferroni corrections were applied across four comparisons. Statistical significance was defined as α = 0.05, and corrected *P*-values are reported.

For track-based measures (5-km run time, heart rate, and cadence), we again assessed normality using the Shapiro-Wilk test. Paired t-tests were used to compare exotendon and natural conditions for each metric. Bonferroni corrections were applied across three comparisons, with α = 0.05, and corrected *P*-values reported.

## Acknowledgements

The authors thank Julie Muccini for assistance with the experiments and the runners who dedicated their time.

## Data availability statement

The code and supporting links to data for the findings of this study are openly available at https://github.com/stingjp/ExotendonPredictions.git.

## Declaration of AI-assisted technologies

During the preparation of this work the authors used OpenAI’s ChatGPT as an editing tool of which the authors manually implemented appropriate changes.

