## Supplemental materials for "Simulation-guided design of exotendons to reduce the energetic cost of running"

### Simulation validation at 2.7 m/s

To validate the simulation framework, we first simulated unassisted (referred to as natural), and exotendon running at 2.7 m/s, a speed we have examined previously in experiments. For the simulated results at 2.7 m/s, we compared the changes in energetic cost (Fig S1). Whole-body changes in metabolic cost (Fig S1a) were slightly underestimated in the current simulation framework compared to experimental data and previous simulation studies (mean error: 3.0%; 95% CI: -0.3 to 6.2%). However, the differences in stance phase versus swing phase gait (Fig. S2b) were aligned with previous results. These comparisons showed that the current pipeline was able to capture the key changes in energetic cost and gave us confidence that the simulation framework is capturing the runner’s adaptation to an exotendon, without tracking any exotendon data.

The simulated kinematics, moments, and ground reaction forces for the natural (Fig S2 orange) and exotendon (Fig S2 purple) simulations closely resembled experimental data at this running speed (Fig S2 shaded regions), giving us confidence that the simulation pipeline would perform well for a faster running speed. The framework reproduced changes in experimental joint kinematics (root-mean-squared-error (RMSE): 4.8 ± 2.2°; Pearson r = 0.97 ± 0.01), and joint moments (RMSE: 0.3 ± 0.1 Nm/kg; r = 0.94 ± 0.02) at the hip, knee, and ankle during running at 2.7 m/s with the previously published exotendon parameters (Fig S2).

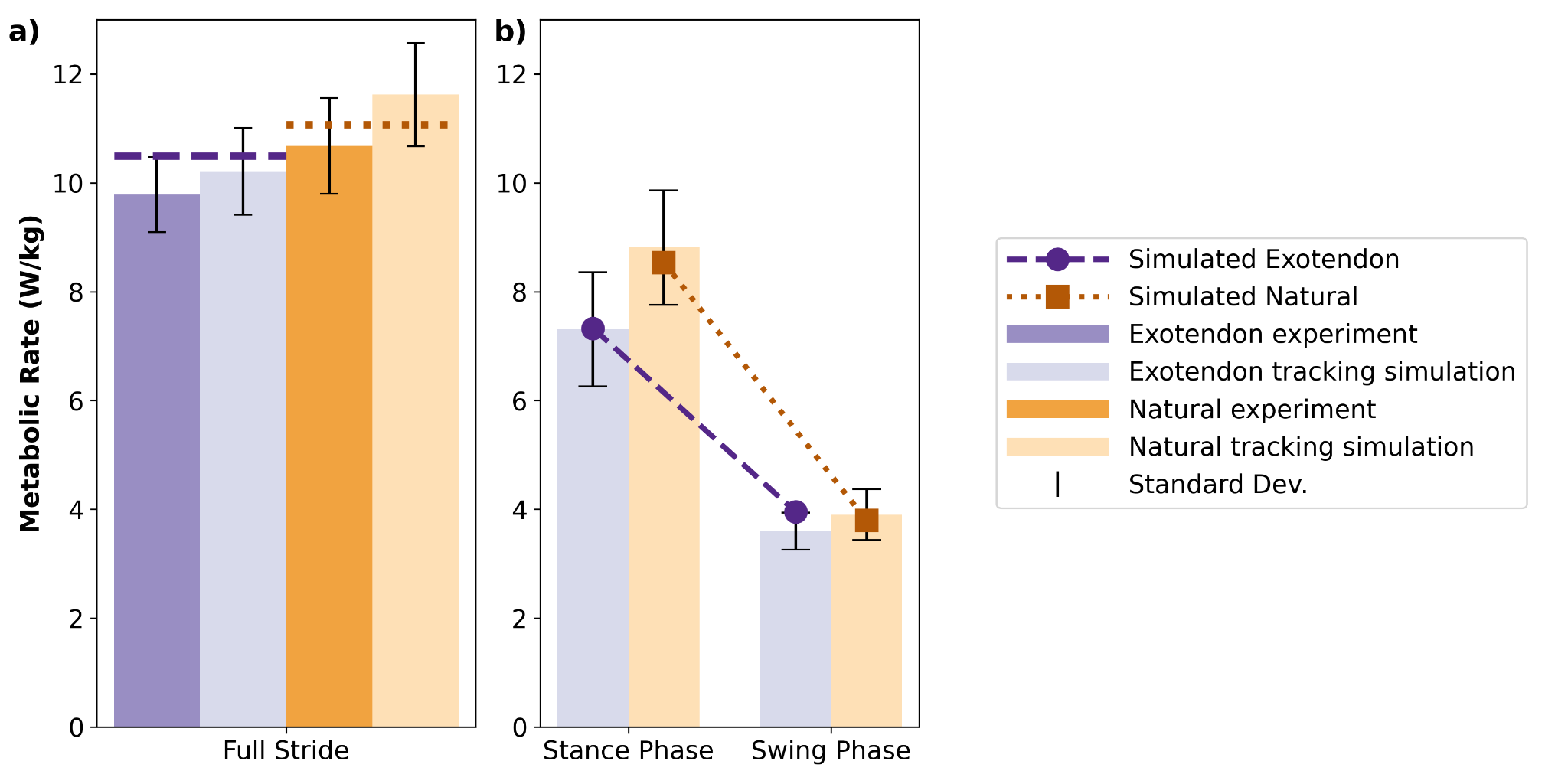

**Fig S1:** The average energetic cost is shown for natural (orange) and exotendon (purple) running simulations. The darkest bars represent the experimental results from Stingel et al. (2023), the light shaded bars represent the simulated data from Stingel et al. (2023), and the dashed and dotted lines highlight results from the current simulations. The error bars on previous studies’ data signify the standard deviation across subjects. a) The average metabolic rate during the full stride is computed for the whole body, while b) stance phase (left) and swing phase (right) are computed for single leg during that segment of the gait cycle.

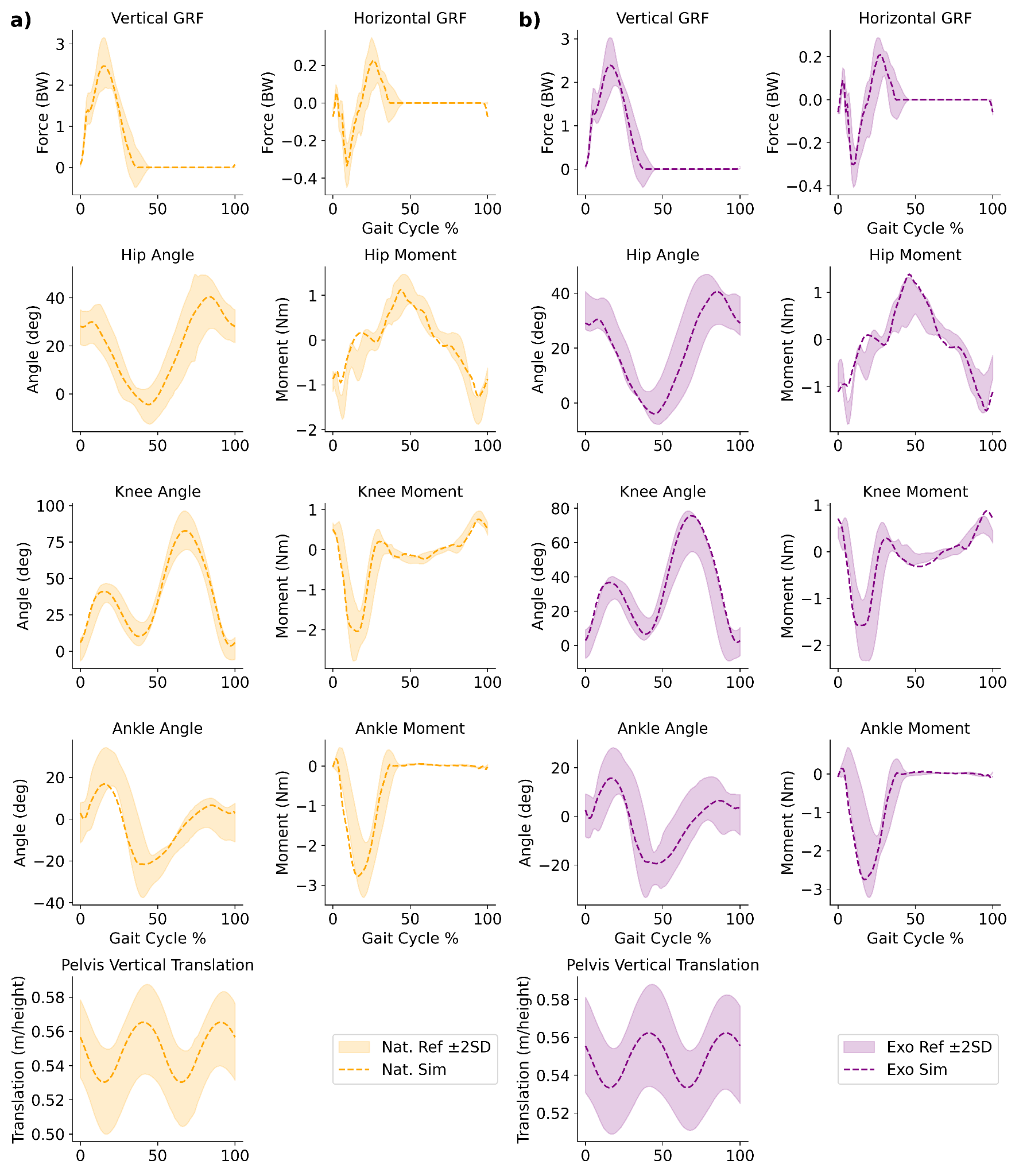

**Fig S2:** Simulated (dashed line) natural (orange) and exotendon (purple) running are plotted in comparison to 2 standard deviations about the mean of previously published reference data (shaded area) of running at 2.7 m/s. A full gait cycle starting with heel strike is shown, with the data representing the right leg where applicable. Vertical and horizontal ground reaction forces are shown (first row). The joint angles and moments are shown for the hip, knee, and ankle (second-fourth row). Finally, the vertical position of the pelvis is also shown.

To explore the ways in which the current simulation framework was adapting to the exotendon, we compared the muscle-level metabolic changes when running at 2.7 m/s to those of prior simulation studies with a more detailed musculoskeletal model (13) (Fig S3). The change in muscle-level biomechanics across the major muscle groups agrees well between simulated frameworks and models, providing confidence that the current simulation framework was capturing salient features of exotendon gait.

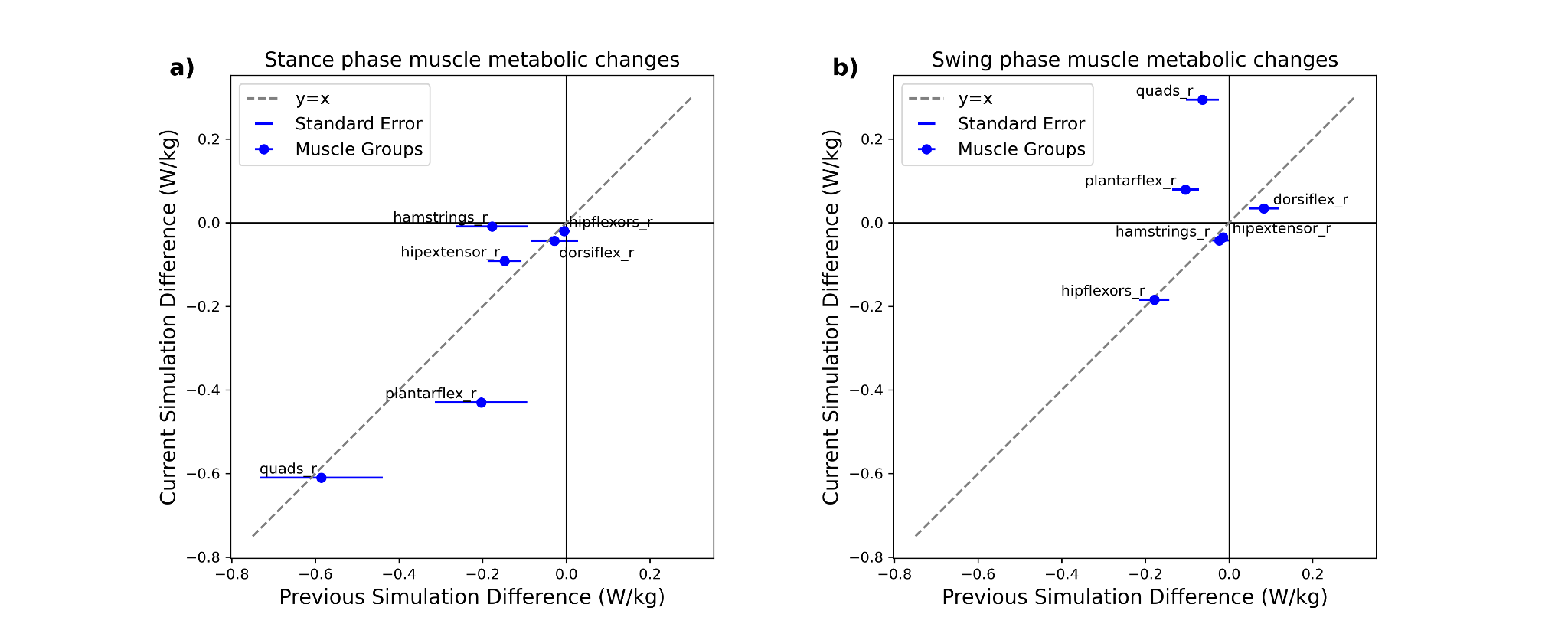

**Fig S3:** *a)* The difference in average metabolic rate during the stance phase of gait for the major muscle groups in the current simulation framework’s musculoskeletal model is plotted with respect to previously published results in Stingel et al. (2023). The previously published results (x-axis) are averaged over 7 subjects, and the error bars designate the standard error across subjects. The current simulation framework’s results (y-axis) signify the energetic cost for a single subject. The muscles are combined into six functional groups including the hip flexors and extensors, hamstrings and quadriceps, and plantar and dorsiflexors. *b)* The same data is shown for the swing phase of the gait cycle.

### Simulation Validation at 4 m/s

To further validate the simulation framework, we compared simulated natural running joint kinematics (RMSE: 8.4 ± 4.2 degrees; *r* = 0.94 ± 0.03) and moments (RMSE: 0.5 ± 0.1 Nm/kg; *r* = 0.91 ± 0.04) at the hip, knee, and ankle at 4 m/s (25) (Fig S4). These results showed that the framework was able to reliably generate simulations of running at 4 m/s, which gave us confidence once again that simulations at this speed will accurately represent reality.

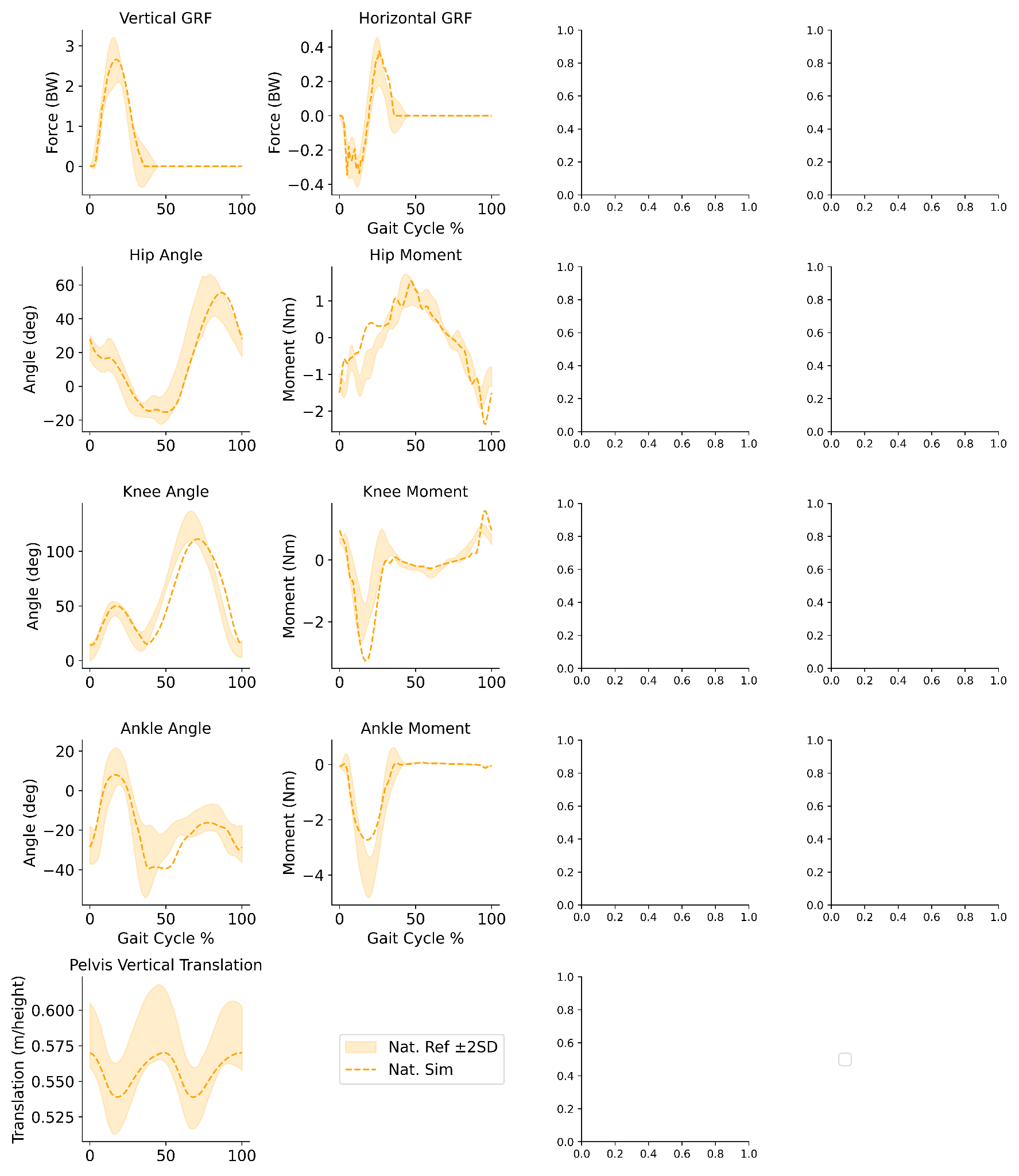

**Fig S4:** Simulated (dashed line) natural (orange) running is plotted in comparison to 2 standard deviations about the mean of previously published reference data (shaded area) of running at 4.0 m/s. A full gait cycle beginning with heel strike is shown, with the data representing the right leg where applicable. Vertical and horizontal ground reaction forces are shown (first row). The joint angles and moments are shown for the hip, knee, and ankle (second, third, and fourth rows, respectively). Finally, the vertical position of the pelvis is shown.

### Running with an exotendon at 4 m/s

We compared the best performing exotendon design’s resulting gait kinematics (Fig S5, S6) and moments (Fig S6) to those of the simulated natural running gait. The largest changes in gait occur at the knee joint. While using an exotendon, the simulated subject utilized a straighter knee, with lower knee extension moment. This change is consistent with the previously studied speed of 2.7 m/s.

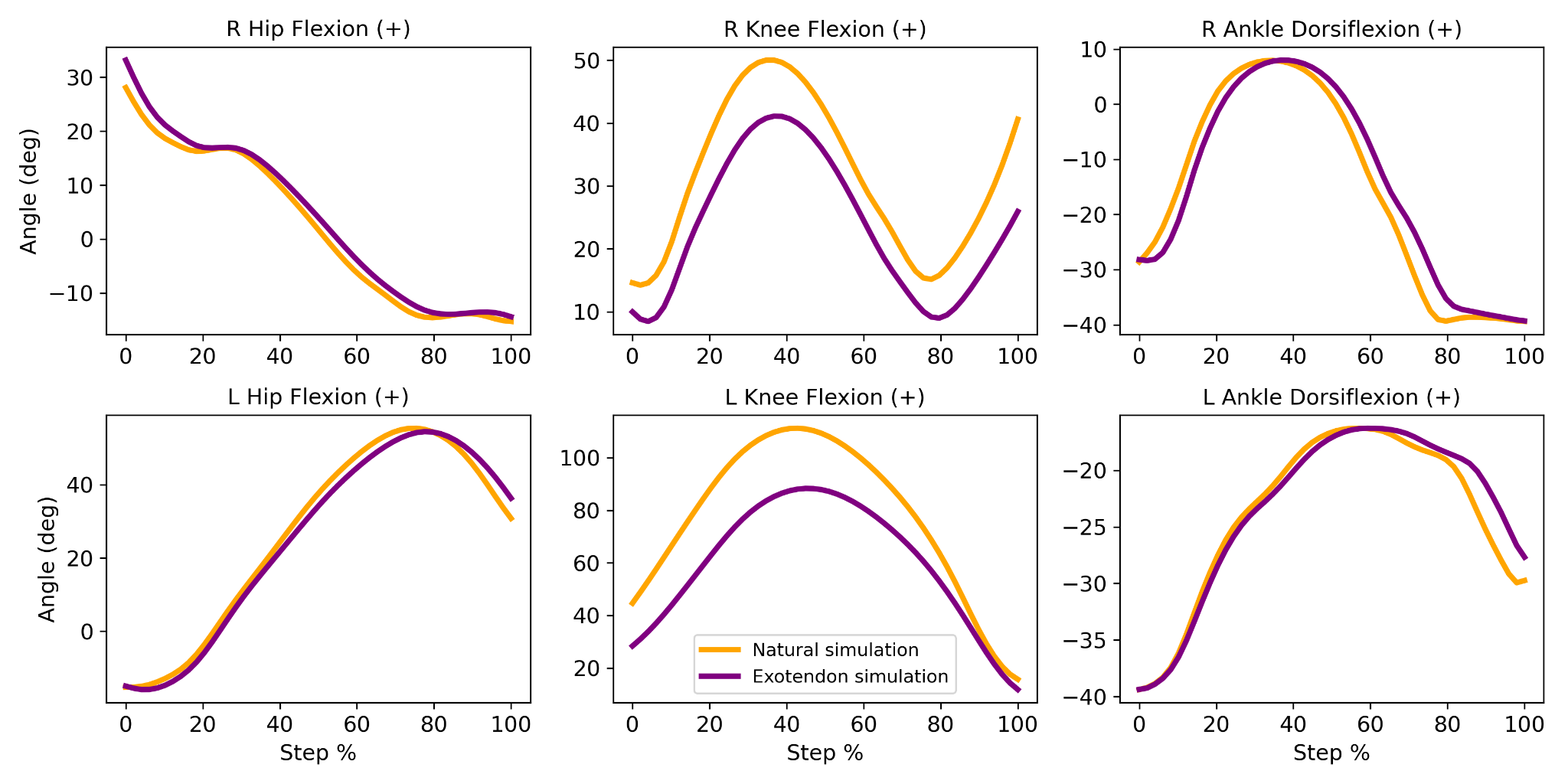

**Fig S5:** Simulated kinematics are shown for natural (orange) and exotendon (purple) running simulations at 4.0 m/s. The right (top) and left leg (bottom) sagittal hip flexion, knee flexion, and ankle dorsiflexion are shown. Positive angles indicate hip flexion, knee flexion, and ankle dorsiflexion. Each step is oriented to begin with heel strike and end in flight phase at half of the gait cycle.

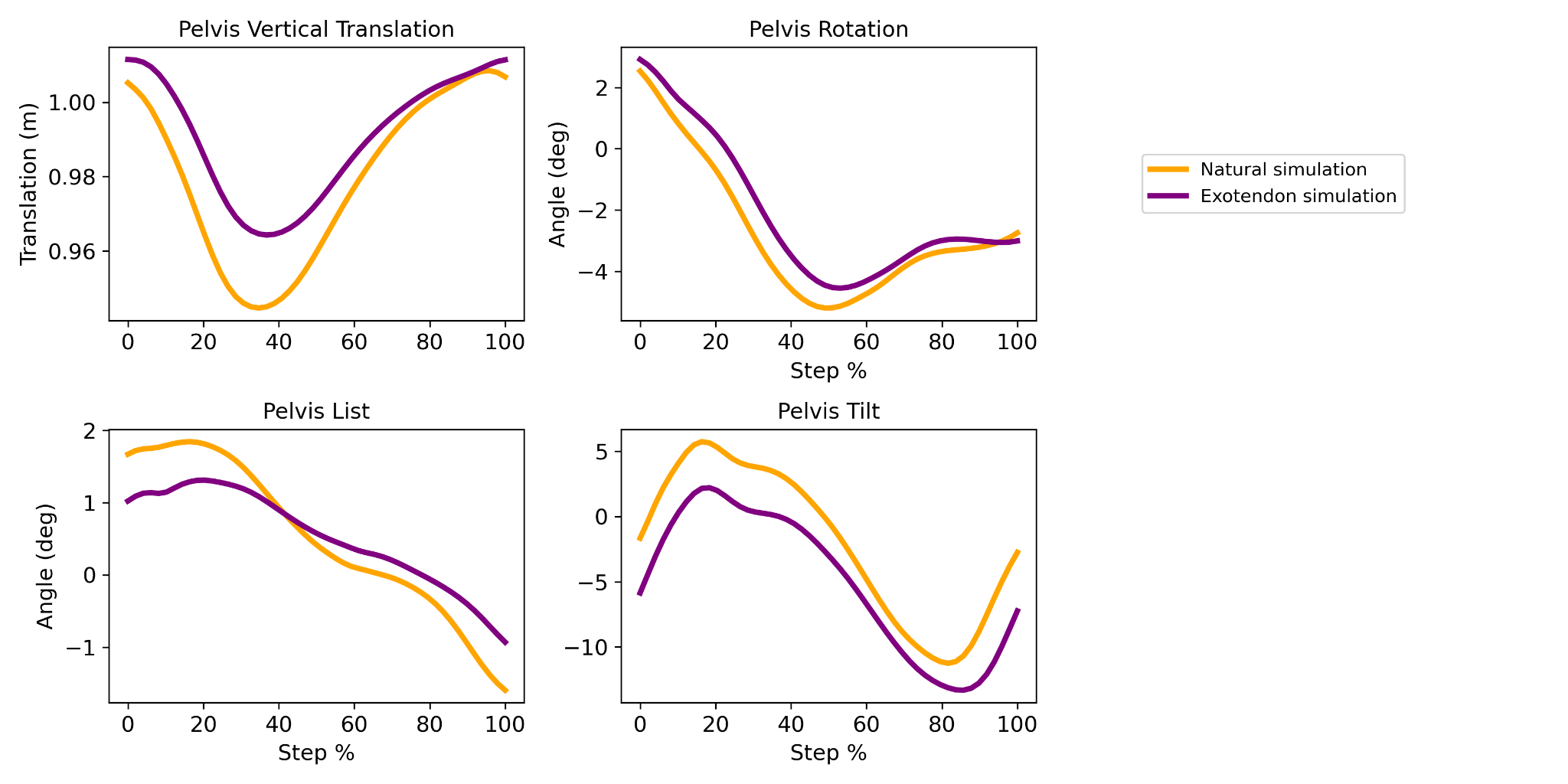

**Fig S6:** Simulated kinematics are shown for natural (orange) and exotendon (purple) running simulations at 4.0 m/s. The pelvis vertical translation (top left), rotation (top right), list (bottom left), and tilt (bottom right) are shown. Each step is oriented to begin with heel strike and end in flight phase at half of the gait cycle.

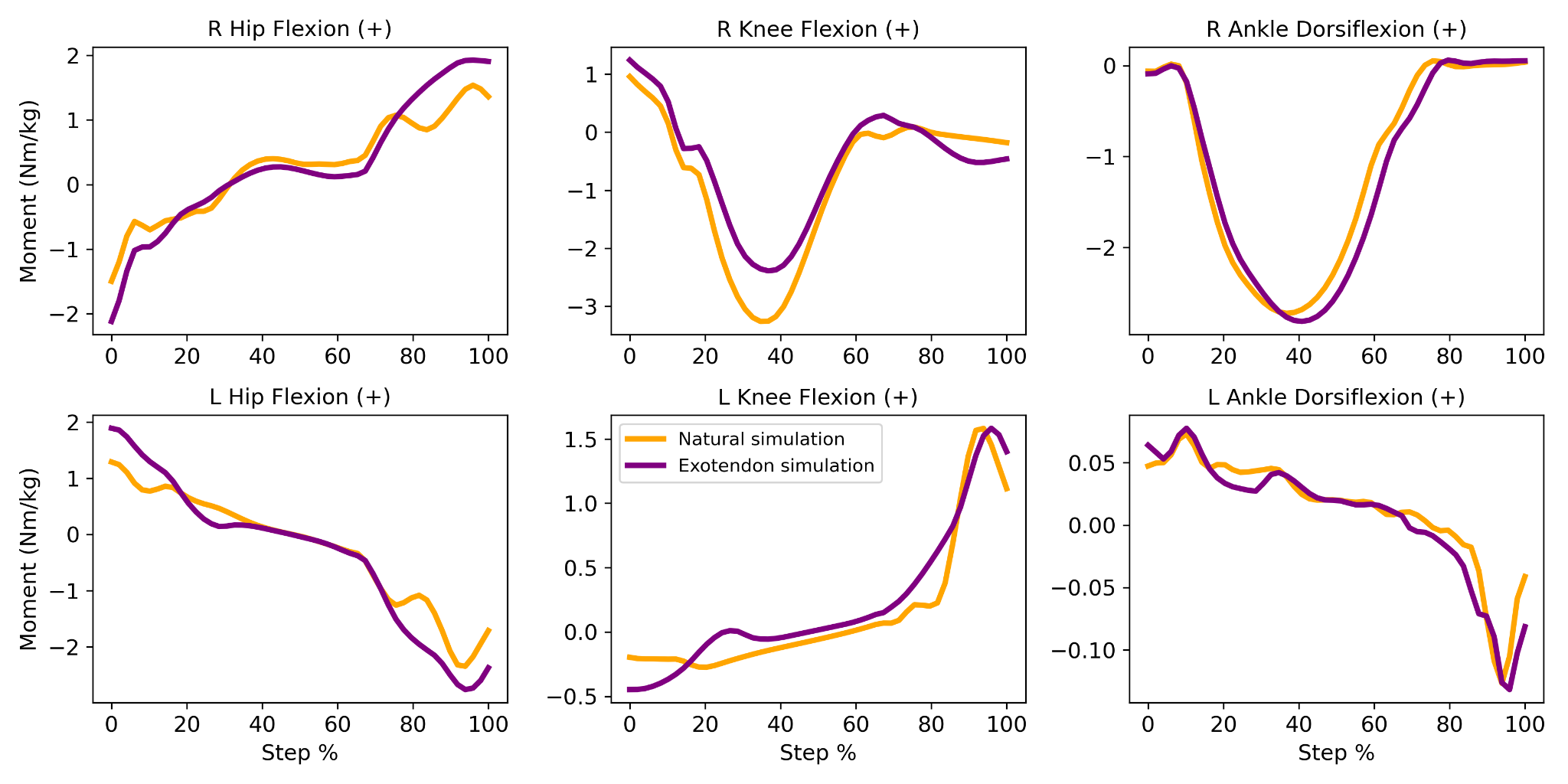

**Fig S7:** Simulated joint moments are shown for natural (orange) and exotendon (purple) running simulations at 4.0 m/s. The right (top) and left leg (bottom) sagittal hip flexion, knee flexion, and ankle dorsiflexion moments are shown. Positive moments indicate hip flexion, knee flexion, and ankle dorsiflexion. Each step is oriented to begin with heel strike and end in flight phase at half of the gait cycle.

The most beneficial exotendon design varied across subjects during the in-lab energetic cost trials (Fig S8). There was at least one runner who achieved their minimum energetic cost with each of the exotendon designs. One participant did not save energy with any of the designs, and there was a total of five other trials across participants where they did not save energy with a given exotendon. The largest number of runners (6) achieved their minimum energetic cost while using the medium-original exotendon; two with the long-compliant exotendon, two with the short-stiff exotendon, and one with the long-stiff exotendon.

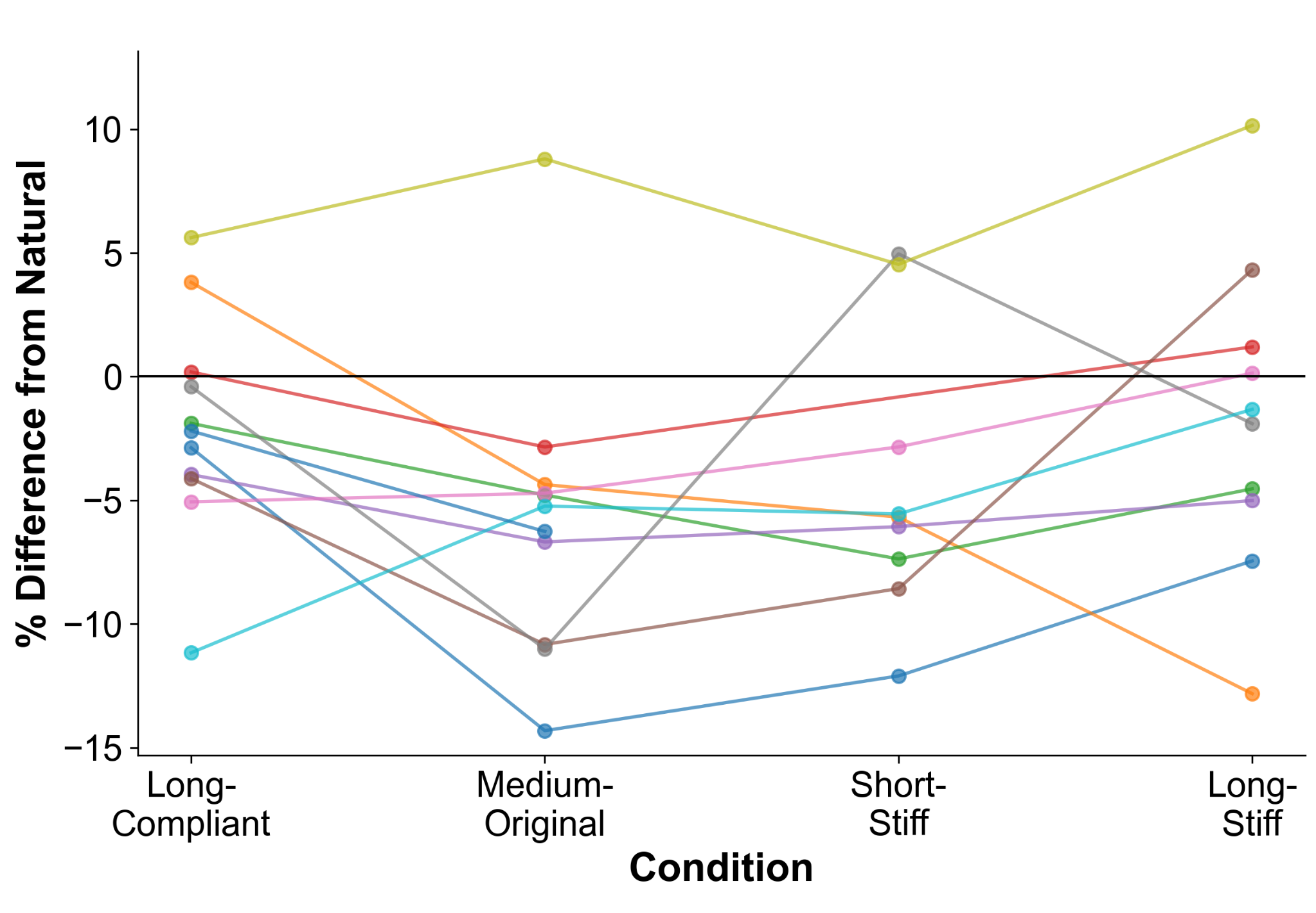

**Fig S8:** Energetic changes from *N*=11 participants running at 4 m/s experimentally. Percent change in energetic cost (net metabolic power) relative to natural running is shown for each participant and exotendon design. Individual participant’s data is connected with a different colored line.

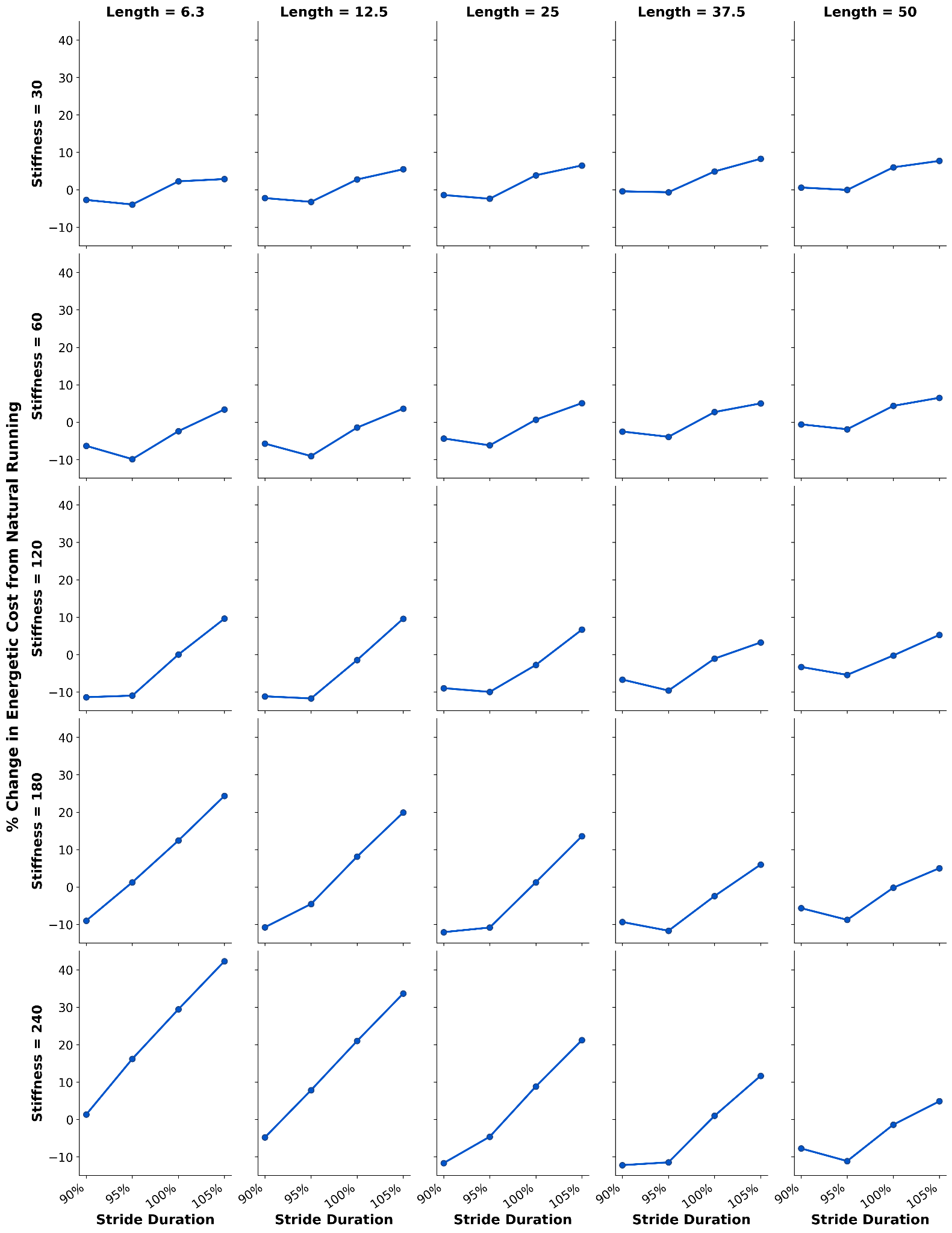

**Fig S9:** Energetic cost across exotendon designs at varying stride durations. Grid showing percent change in energetic cost from natural running for exotendon stiffness (rows: 30–240 N/mm) and length (columns: 6.3–50 % leg length) combinations. Each subplot displays four measurements corresponding to stride durations of 90%, 95%, 100%, and 105% of natural baseline stride duration. Negative values indicate metabolic savings; positive values indicate increased cost relative to natural running.
